# A single-cell catalogue of regulatory states in the ageing *Drosophila* brain

**DOI:** 10.1101/237420

**Authors:** Kristofer Davie, Jasper Janssens, Duygu Koldere, Uli Pech, Sara Aibar, Maxime De Waegeneer, Samira Makhzami, Valerie Christiaens, Carmen Bravo González-Blas, Gert Hulselmans, Katina I. Spanier, Thomas Moerman, Bram Vanspauwen, Jeroen Lammertyn, Bernard Thienpont, Sha Liu, Patrik Verstreken, Stein Aerts

## Abstract

The diversity of cell types and regulatory states in the brain, and how these change during ageing, remains largely unknown. Here, we present a single-cell transcriptome catalogue of the entire adult *Drosophila melanogaster* brain sampled across its lifespan. Both neurons and glia age through a process of “regulatory erosion”, characterized by a strong decline of RNA content, and accompanied by increasing transcriptional and chromatin noise. We identify more than 50 cell types by specific transcription factors and their downstream gene regulatory networks. In addition to neurotransmitter types and neuroblast lineages, we find a novel neuronal cell state driven by *datilografo* and *prospero*. This state relates to neuronal birth order, the metabolic profile, and the activity of a neuron. Our single-cell brain catalogue reveals extensive regulatory heterogeneity linked to ageing and brain function and will serve as a reference for future studies of genetic variation and disease mutations.

## Introduction

Brain function depends on a complex network of specialized neuronal and glial cell types. Single-cell technologies allow us to map this diverse cellular landscape, and single-cell RNA-seq in specific parts of both the human and mouse brains have already revealed several unique cell types (Lake et al., 2016; Tasic et al., 2016; Zeisel et al., 2015). However, a compendium of all cell types in a complete brain at specific time points during ageing has not yet been reported. This is important because cell compositions may change during ageing, and the timing of normal and pathological cell loss, and the identity of these cellular subtypes remain poorly described. Comprehensive and unbiased brain-wide single-cell sequencing will facilitate understanding of the cellular basis of a brain and provide insights on the gradual loss of fitness and cognition in ageing, and their connections to neurodegeneration and inflammation (Tulving and Craik, 2005; Wyss-Coray, 2016).

The *Drosophila* brain is ideal to build a catalogue of cell types of a complex functioning brain because it encodes an extensive array of intricate behaviours (Owald et al., 2015), while only consisting of approximately 100,000 cells, of which the majority (85-90%) are neurons (Ito et al., 2013; Kremer et al., 2017). In addition, the fly brain has been well described in terms of neuronal connectivity (Chiang et al., 2011) and in terms of connecting behavioural traits to neuronal subpopulations (Robie et al., 2017). Building a reference map of cell types, at singlecell resolution, will be useful for future studies, for example to compare single-cell profiles related to natural and evolutionary variation, and related to disease-associated mutations.

Cell types and cell states are encoded in the genome as “regulatory programs”, and maintained by gene regulatory networks, in which combinations of transcription factors regulate the expression of their target genes. By sequencing the transcriptome of individual cells, we can now measure the output of the regulatory system without the loss of information caused by the signal averaging from bulk technologies. We recently showed that transcription factor subnetworks that are active in specific cell types can be predicted by combining single-cell RNA-seq with the *cis*-regulatory sequence information contained within the enhancers of regulated genes (Aibar et al., 2017). Understanding the regulatory mechanisms underlying brain cell types might be key to disentangling different diseases of the brain.

Here we build a comprehensive catalogue of the cell types in the adult *Drosophila* brain, characterize the regulatory networks underlying these cell types, link regulatory states with metabolic potential and excitability, and map cell state changes that occur during ageing in a complex brain.

## Results

### Single-cell RNA-seq of the adult brain

We used droplet microfluidics, a high-throughput and scalable methodology to achieve a high cell-coverage of the entire *D. melanogaster* brain. In a pilot study, we assessed the sensitivity of two droplet based scRNA-seq approaches: Drop-seq (Macosko et al., 2015), and the 10x Chromium (10x Genomics); each applied to dissociated adult brains of newly-eclosed adults. We consistently detected the highest number of genes per cell on the 10x Chromium platform: 1,468 genes per cell, compared to 250-400 genes per cell for Drop-seq (Figure S1). We continued with the 10x Chromium in this study.

To take genetic diversity between domesticated *D. melanogaster* strains into account we dissected brains from two different lab strains, one that has been domesticated recently (DGRP-551) and one that has been domesticated more than 70 years ago (w^1118^). We used animals precisely aged to eight different time points, ranging from newly-eclosed to 50-days-old (see Methods), covering 26 biological replicates in total (Table S1, Table S2). We achieved a median sequencing depth of 53,553 sequence reads per cell, with median sequence saturation rate of 81.5 %. The total number of distinct genes detected across the dataset was 12,436 protein-coding genes and 2,164 non-coding RNAs. Interestingly, this is significantly higher than the total number of genes expressed in *Drosophila* embryo cells (~8,000) (Karaiskos et al., 2017), and could be suggested to be due to the higher diversity of cell types in the brain compared to the embryo.

To provide an overview of the cell type diversity in the fly brain, we utilized the entire dataset and generated t-Distributed Stochastic Neighbor Embedding (t-SNE) plots in SCENIC (Aibar et al., 2017) (Figure 1A) and Seurat (Satija et al., 2015) (Figure S2). We also generated individual plots for each of the two domesticated lines and found a highly similar structure, robust to t-SNE parameter choice (Figure 1B/C). These unsupervised clustering methods distinguish multiple clusters of neurons and glia, identified by the expression of the neuronal marker *elav* and the glial marker *repo* (Figure 1D). We identified a relatively low percentage of all sequenced cells as glia (6.4%), while the fly brain consists of about 15% glial cells (Kremer et al., 2017). However, this is because we detected a lower number of mRNAs and distinct genes in glial cells than in neurons, causing a larger fraction of glia to not pass our filtering thresholds. These filters were based on a combination of Cell Ranger (10x Genomics) and scater (McCarthy et al., 2017a) suites (see Methods). Lowering the filter stringency, we retained more than 129,000 cells (Figure S3A) and the fraction of glia increased to 11.5% (Figure S3B/C). Furthermore, we verified differential expression of glial and neuronal markers other than *repo* and *elav*. Glial cells in our dataset express typical glial genes such as *Gs2, nrv2*, and *Eaat1* (all among the top 20 most up-regulated genes in glia), while neurons express typical neuronal genes, such as *nSyb* and *Syt1* (in the top 20 most up-regulated genes in neurons). Interestingly, the most abundant and most differentially expressed genes between neurons and glia are long non-coding RNAs (lncRNA), such as *noe* and *CR31451* in neurons, and *MRE16* in glia (Figure 1E). These very highly expressed genes subsequently enabled the identification of additional neurons and glia in the cells that were initially filtered out. This strategy again revealed that a large fraction of the omitted cells were indeed glia (Figure S3B/C).

**Figure 1.**
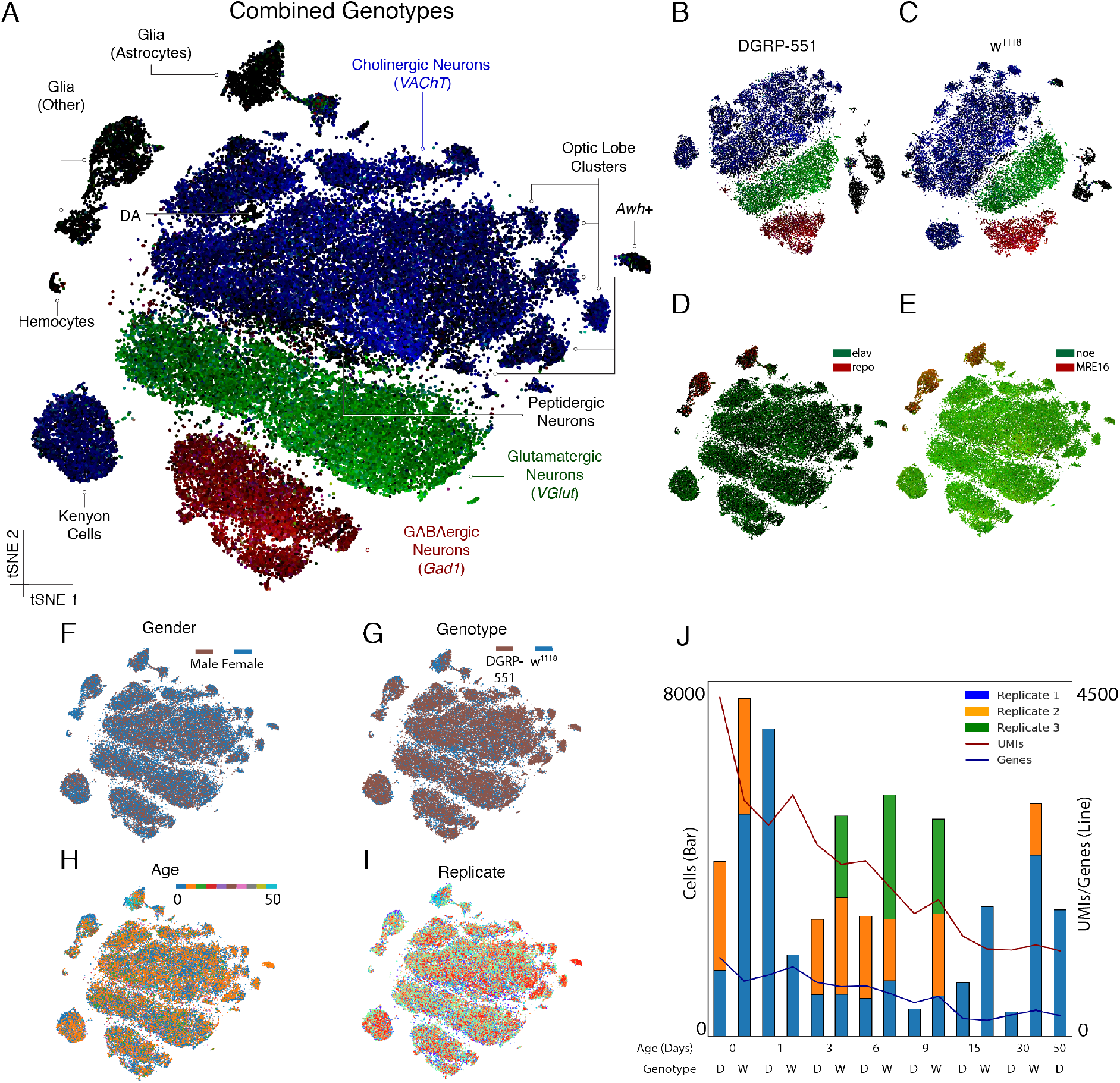
Single-cell RNA-seq of the adult brain. (A) t-SNE plots generated from SCENIC analysis, combining the two genotypes, cluster by presynaptic neurotransmitter type. Cells are colored by expression of *VAChT* (blue), *VGlut* (green) and *Gad1* (red). (DA: Dopaminergic neurons) (B-C) Separate t-SNEs for the lines DGRP-551 and w^1118^, show highly reproducible cell types and clustering. (D) Neurons and glia are distinguished by the neuronal marker *elav* (green) and glial marker *repo* (red) as well as (E) Neuronal and glial specific lncRNAs *noe* (green) and *MRE16* (red). (F-I) No bias is observed in the clustering for gender, genotype, age, and replicate respectively. (J) Numbers of single cells per 10x Chromium run, as well as median number of UMIs and genes detected, which decline by age (D: DGRP-551; W: w^1118^). See also Figure S2 and Figure S3.

Our starting material consisted of equal amounts of male and female fly brains and this was very well recapitulated in our catalogue. We used expression of male-specific lncRNAs regulating dosage compensation, *roX1* and *roX2* (Figure 1F) and found 49% male cells and 51% female cells. We also confirmed that both genotypes (Figure 1G), all ages (Figure 1H), and all replicates (Figure 1I) mix well within our clustering. These results indicate that we equally sampled cells from all brains entering our analyses, and that our resource encompasses a high-quality dataset of the expression profiles of 56,902 cells (Figure 1J) of both neuronal and glial subtypes in the male and female *Drosophila* brain.

### Ageing is accompanied by regulatory erosion

Our dataset contains cells from brains of different ages, from newly-eclosed (0 days) to old (50 days). This allowed us to follow cell state changes by examining gene expression in different cell types during the ageing process. Strikingly, the total RNA content per cell, as measured by the number of unique molecular identifiers (UMI) decreased exponentially (R^2^ = 0.907 for DGRP-551, and 0.840 for w^1118^) during the first nine days of adult life, with a highly similar trend for the two strains (Figure 2A). As expected, this global decline of UMIs coincided with a drastic decrease in number of genes expressed per cell (Figure 2B). This decline of RNA content is in agreement with earlier observations of total RNA decline during the first ten days of adult life of *D. melanogaster* (Tahoe et al., 2004). Some proteins, such as the dopaminergic neuron-specific protein Pale, a tyrosine hydroxylase, also significantly decrease between old and young flies, as demonstrated by immunofluorescence (Figure S4).

**Figure 2.**
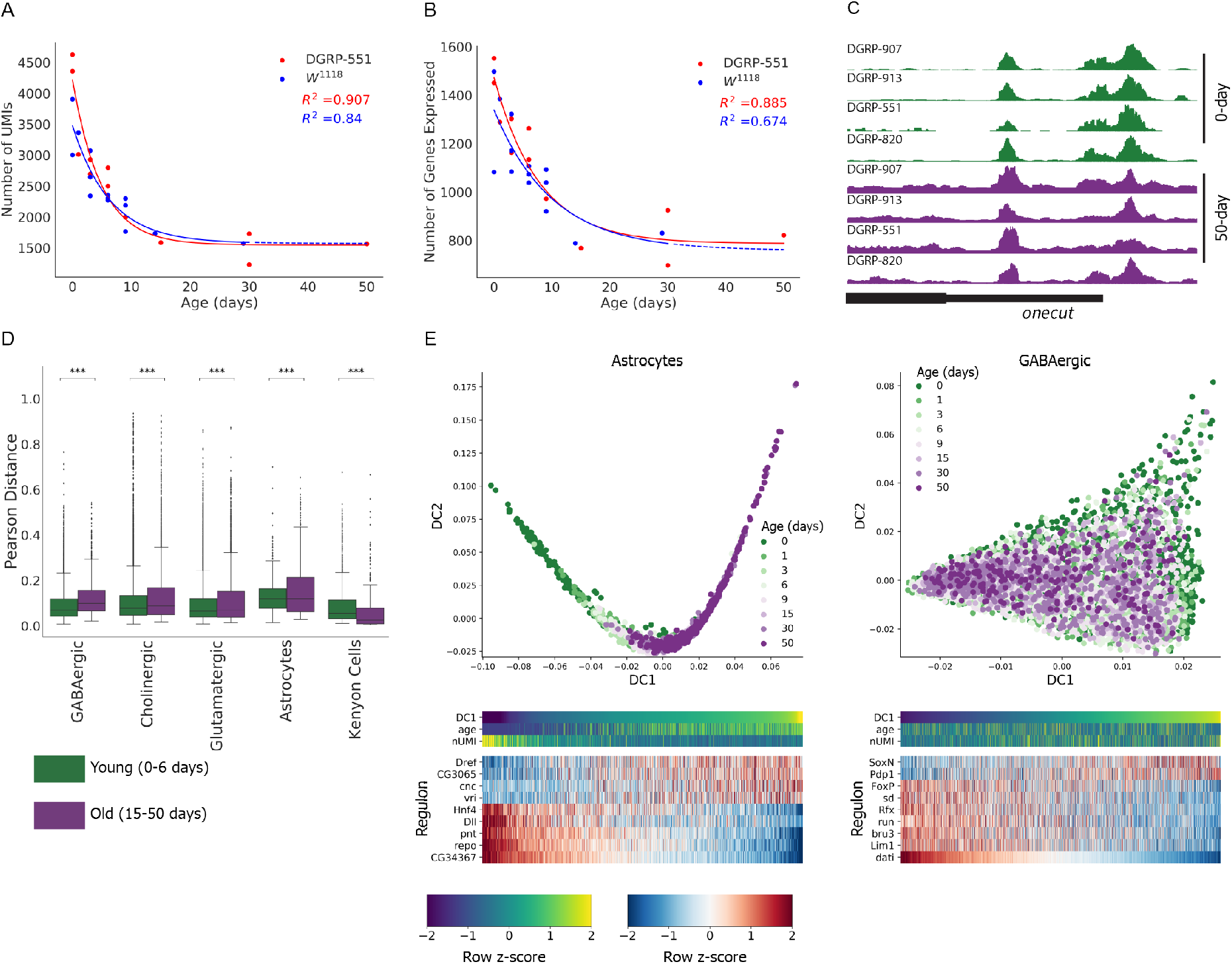
Age-dependent regulatory erosion. (A-B) The number of UMIs (A) and genes (B) declines exponentially with age, following highly similar trends in both DGRP-551 and w^1118^ genotypes. (C) In young brains (0 days), the distribution of open and closed chromatin regions (measured with ATAC-seq) is reproducible across cell- and genotypes, whereas old brains (>50 days) have significantly higher noise in nucleosome positioning. The four replicates are four different DGRP lines. (D) The noise in gene expression (measured as Pearson distance) increases significantly with age across all major cell types, except the Kenyon cells. (E) Astrocytes, but not neurons can be sorted in a diffusion trajectory that correlates with age. In neurons (only GABAergic shown), the first diffusion component correlates strongest with the expression of *dati*. See also Figure S4.

If the abundance of all proteins decreases with age, including transcription factors and histones, we expected to observe changes in the epigenome of old brains. To test this idea, we utilized ATAC-seq, a methodology to map genome-wide accessible chromatin (Buenrostro et al., 2013). We observed significantly higher background (i.e., more noise) in the ATAC-seq signal of 50-day-old DGRP-551 flies compared to the signals in 0-day old flies (Figure 2C) (p=0.0159, paired t-test). This was not strain-specific as we recapitulated these findings in three other *D. melanogaster* strains (Figure 2C). These data indicate that cells in older brains either contain fewer nucleosomes, and/or wrongly positioned nucleosomes, hence the term *“regulatory erosion”*. Note that the chromatin signal in accessible regions remained present as peaks (Figure 2C), therefore the increase of signal-to-noise was not due to a decrease in the signal (i.e., the ATAC-seq peaks), thus favoring the hypothesis of an overall decrease in compaction rather than an increase. As a consequence of this regulatory erosion, transcriptional control was affected, as seen by the increased noise of gene expression for all major cell types, except the Kenyon cells (Figure 2D).

To investigate whether ageing occurs through specific cell state changes along a continuous trajectory, we used diffusion maps (Angerer et al., 2016) to place the cells in a pseudo-temporal order (Figure 2E). The trajectories obtained from astrocytes followed the ageing process, with a decrease in expression of glial markers (e.g., *pnt* and *repo*). In contrast, neuronal cell types were more stable over age, even when the number of UMIs dropped drastically (Figure 2A). In neurons, other factors drive expression variation along the first diffusion component. Particularly, for the three main neurotransmitter types, expression of *dati* appeared to be a key driver of cellular diversity, independent of ageing. We describe the patterns of *dati* expression in detail further below. In conclusion, ageing in *Drosophila* has profound and widespread erosion effects on transcription and chromatin in general, and specific gene expression effects in glia. Neuronal states seem to be steadier during ageing, since not age, but specific neuronal identities underlie most of the variance observed in unsupervised clustering and trajectory inference.

### Principal neuronal cell types represent neurotransmitter types and neuroblast lineages

Single-cell clustering for both genotypes, either using Seurat or SCENIC, primarily identified the neuronal subtypes based on their presynaptic specialization, thus the neurotransmitter they produce and release (Figure 1A, Figure S2). These clusters were identified using known markers, particularly, genes encoding neurotransmitter biosynthetic enzymes, neurotransmitter receptors, neuropeptides, and neuropeptide receptors (Crocker et al., 2016) (Figure 3A). The most prominent cluster in our dataset consists of cholinergic neurons (*VAChT* and *ChAT* expression) with 24,802 of the 56,902 cells (43.6%) (both lines combined), followed by 13,296 glutamatergic neurons (*VGlut*), 6,177 GABAergic neurons (*Gad1*), and 2,998 Kenyon cells (*ey, prt*). Smaller populations of 348 dopaminergic (*ple & Vmat*), 315 octopaminergic (*Tdc2 & Vmat*) and 256 serotonergic neurons (*SerT & Trh*) are also identified by their expression of specific markers (Figure 3A). We further identified a set of 137 neuroendocrine cells that express *dimm* (Park et al., 2008); 41 of these being *Pdf*-positive clock neurons, and at least 16 insulin-producing cells marked by expression of *Ilp2* and *Ilp5*. Thus, we recapitulate prior knowledge, although our dataset now delivers specific gene signatures and markers for each of these clusters.

**Figure 3.**
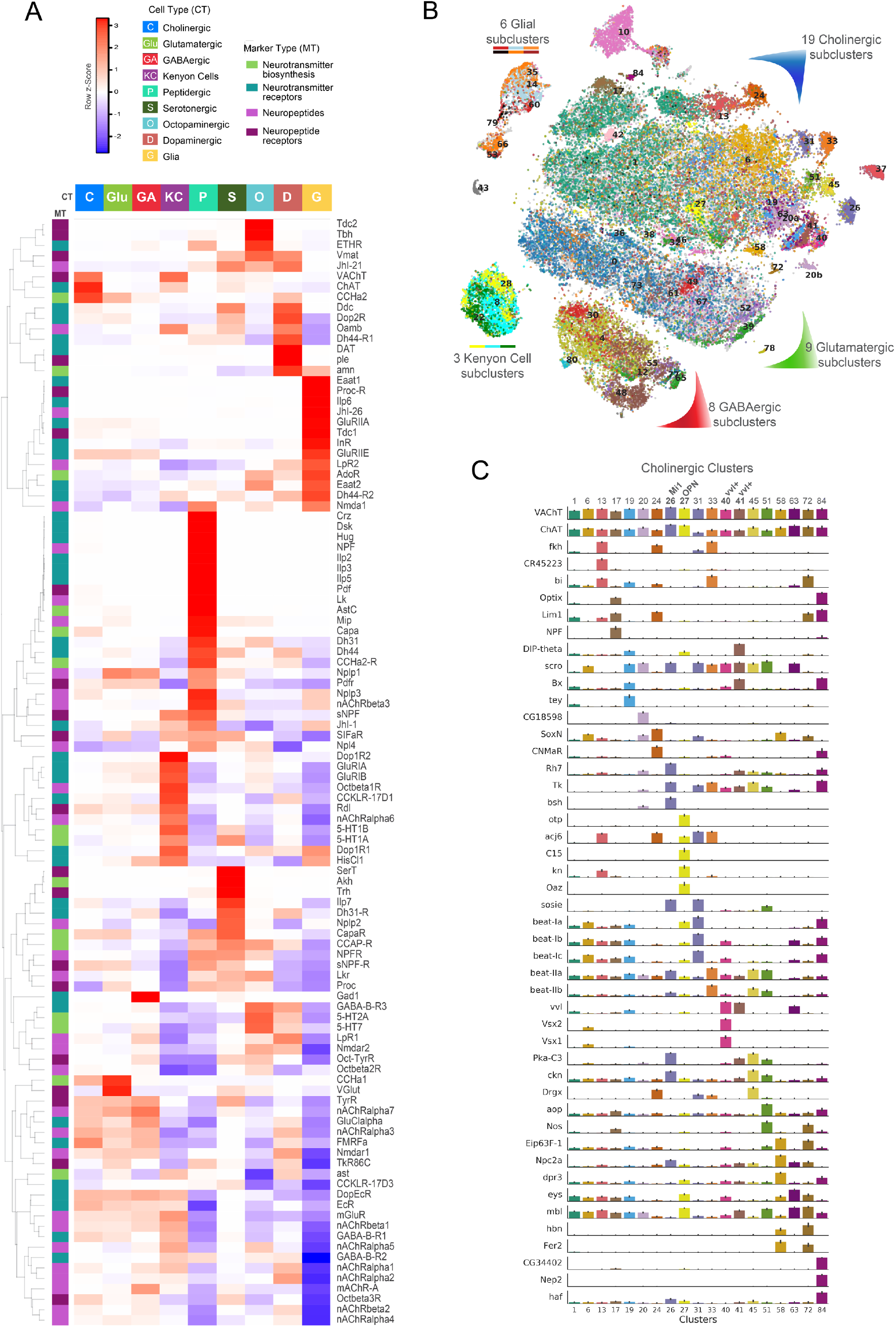
Diversity of neuronal and glial cell types in the brain. (A) Heatmap of the main neuronal and glial clusters (cfr. Figure 1), with expression of neurotransmitter biosynthetic enzymes, neurotransmitter receptors, neuropeptides, and neuropeptide receptors (Crocker et al., 2016). (B). SCENIC t-SNE colored by matching Seurat clusters, highlighting the diversity of neuronal subtypes. The largest number of discrete neuronal subtypes are found for cholinergic neurons. (C) Marker genes for each of the cholinergic subclusters identified by Seurat. See also Figure S5.

Next, we estimated the coverage of cell types contained in our dataset. We compared the numbers of single cells in the specific clusters we identified to previously estimated numbers of cells of a specific type. Our dataset identified 2,998 Kenyon cells and 348 dopaminergic neurons, while a single brain is estimated to contain around 4,000 Kenyon cells and 280 dopaminergic neurons (i.e. 75% and 100%). Furthermore, a single brain contains 16 Pdf neurons whereas we found 41 Pdf neurons (>100%). Based on these proportions, we have sampled the resident cell types in the fly brain between 0.7-1x, suggesting we have identified most of the neuronal subtypes in the fly brain. Of the known glial subtypes, we identified the surface glia, cortex glia and astrocyte-like glia (Kremer et al., 2017; Omoto et al., 2016), as illustrated by marker gene expression and t-SNE clustering (Figure S5). However, the glial subtypes show more overlap of marker gene expression than neuronal subtypes, suggesting more overlap of glial cell states. Finally, a group (174 cells) of hemocytes (expression of *Hml*) were detected, as well as 146 cells that resemble photoreceptors (expression of *gl* and *Pph13*), likely due to incomplete removal of the retina during the dissection.

Furthermore, both SCENIC and Seurat detected multiple neuronal subclusters within the main neurotransmitter classes, each with a specific set of marker genes (Figure 3B/C). Within the cholinergic neuron cluster, we detected 19 smaller subclusters which can be further identified. In *Drosophila*, the major neuronal subtypes are determined during development from neuroblast lineages, of which several have been fate mapped to adult neurons (Li et al., 2013a). Some of the detected cholinergic optic lobe subtypes correspond directly to these neuroblast lineages. For example, we identified the Mi1 lineage by co-expression of the specific transcription factor *brain specific homeobox (bsh)*,the non-specific factor *homothorax (hth)* (Li et al., 2013b). We also detect a cluster of *drifter/vvl* positive neurons in the optic lobe as previously reported (Komiyama et al., 2003). Additionally, some cholinergic subclusters can be traced back to the central brain with specific markers. For instance, recently *C15, otp, kn* and *acj6* were identified as markers for the olfactory projection neurons (OPN) by single-cell RNA-seq (Li et al., 2017). When testing for co-localization of these markers, we found a matching cluster of OPNs in our data (cluster 27 in Figure 3B/C). Several new clusters feature among the cholinergic neurons, which we could not yet link with spatial or lineage information, and for which we list marker genes in our online resource, to be used in future studies. Here, we further investigate and characterize these clusters using gene regulatory networks.

### Gene regulatory network mapping reveals key transcription factors of neuronal subtypes

Next, we aimed to functionally characterize the candidate cell types described above, namely by mapping their underlying gene regulatory networks. To this end, we analyzed the output of SCENIC, a computational algorithm that infers transcription factor centered co-expression networks through *cis*-regulatory sequence analysis. As SCENIC was only available for human and mouse, we first built the motif databases that allows its application to Drosophila (See Methods). From all 708 *D. melanogaster* transcription factors (TFs), SCENIC identified 150 TFs with their recognition motif significantly enriched in their co-expressed gene sets. Interestingly, we found that most of the 150 co-expression networks (termed ‘regulons’) marked specific clusters of cells (Figure 4A). Several regulons appeared active only in glial cells: we identified the previously known Repo and Pnt regulons to be active (Klaes et al., 1994; Xiong et al., 1994) (although Pnt is also active in hemocytes), but also new regulons such as the Hnf4 regulon.

**Figure 4.**
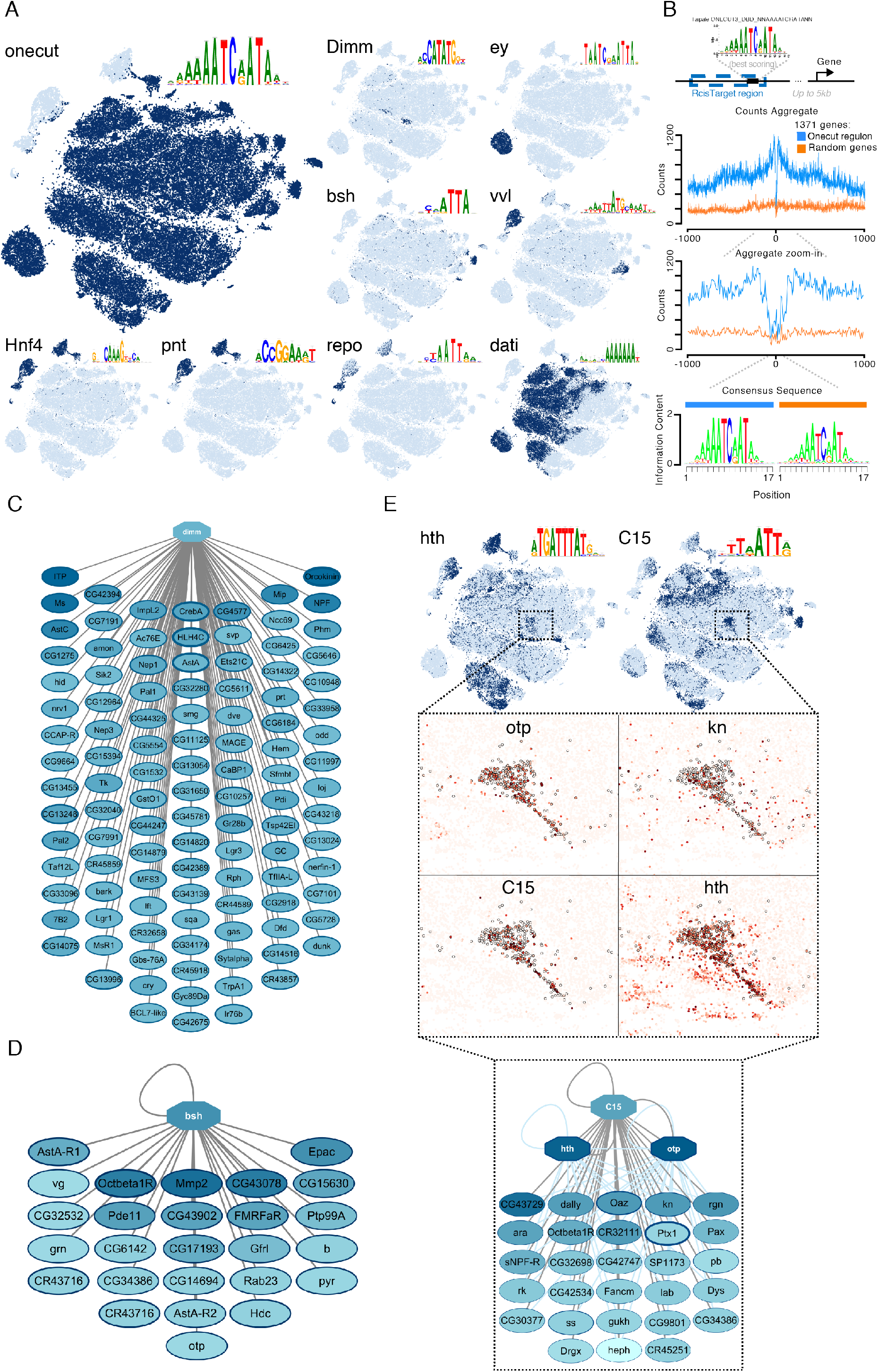
Gene regulatory networks underlie neuronal and glial cell types. (A) Selected regulons from SCENIC, showing cells having that regulon active, and the best enriched motif for that factor. (B) DNA footprint of the Onecut binding site predictions, using ATAC-seq. In blue ATAC-seq signal of the best Onecut site near genes of the Onecut regulon; in orange, the best Onecut sites near randomly selected genes. In Figure S6A we also compare this with background Tn5 insertions. (C) The Dimmed regulon, active in peptidergic neurons. (D) The Bsh regulon, active in the Mi1 lineage of the optic lobe. (E) The C15 regulon, active in the olfactory projection neurons with the expression of four target genes as inset. Otp and Hth were added as co-regulatory factors because their motif is also enriched in the C15 regulon, through an analysis with iRegulon (Janky et al., 2014). We also compare this with background Tn5 insertions. See also Figure S6.

In neurons, pan-neuronal TFs such as Onecut and Pdp1 (and others) appear active. Pdp1 is a Tef homolog, which was found as pan-neuronal factor in the mouse brain using SCENIC with similar motifs enriched for the mouse factor and the Drosophila factor (Aibar et al., 2017). Onecut is a Cux1/2 homolog and its regulon contains 1,371 predicted target genes. These genes are significantly enriched for the Gene Ontology (GO) term “nervous system development” (GO:0007399; FDR 1.21x10^−58^). We also checked if predicted Onecut binding sites reside in open chromatin in the genome. Across four replicates, our data identified Onecut sites to be strongly enriched for ATAC-seq signal (in the DGRP-551 line) (Figure 4B). Importantly, this analysis identified a significant DNA footprint (i.e., protection against Tn5 integration) precisely overlapping the predicted Onecut motif (Figure 4B). We confirmed that this signal was specific, and not due to background Tn5 specificity, by comparing high-scoring Onecut motif instances near random genes (Figure 4B), and by using a recently established model to estimate the number of aspecific Tn5 insertions (Yardımcı et al., 2014) (Figure S6A). Bulk ATAC-seq performed best for the pan-neuronal factors because a large fraction of cells has pan-neuronal enhancers active. However, at least for Onecut, the predicted binding sites were effectively bound in vivo in the brain at the predicted motif. These data independently confirmed that SCENIC predictions are built from bona fide regulatory interactions.

Next, we inferred TF networks for specific neuronal subtypes. For example, we identified Mi1 neurons as being under the control of Bsh. We represent these regulons as networks, where target gene predictions are ranked by their differential expression and genome-wide ranking based on the specific TF motif (Figure 4C-E). In peptidergic neurons, we identified a prominent network regulated by Dimmed. Interestingly, the 111 predicted Dimmed target genes are significantly enriched for the Gene Ontology term “neuropeptide signaling pathway” (GO:0007218; FDR 1.60x10^−9^), with strong target gene co-expression of *7B2, ITP, Ms, Mip* and others (Figure S6B). Finally, SCENIC also identified a regulon defining OPN neurons under the control of C15 (Figure 4E). Among the predicted C15 target genes are *otp* and *kn*, consistent with previous scRNA-seq data of this isolated population (Li et al., 2017) (Figure 3B/C).

To enrich our analyses with additional transcription factors, we screened the markers found by Seurat. From all TFs, we identified 205 as significant markers of cellular subtypes, of which 104 correspond to SCENIC regulons (Figure S6C), illustrating that Seurat and SCENIC are complementary approaches. As an example, the peptidergic cluster was identified using the Dimmed regulon using SCENIC, but was not found by Seurat clustering. Conversely, *mamo* and other TFs were found as marker TFs by Seurat but not as regulons in SCENIC (Figure S6C). The inability to identify regulons in these cases is likely because of the lack of a specific motif for those TFs, or the weak enrichment of a motif near the co-expressed target genes. Finally, a small number of known TFs with specific regulons were not detected, such as *atonal*, expressed in the dorsal cluster neurons (DCN) (Hassan et al., 2000), this may be because of their low abundance. Of note, this does not imply that DCN neurons are not represented in our dataset. Indeed, several distinct *acj6*-positive clusters were identified, one of which likely represents the DCN. In conclusion, analysis of our dataset using SCENIC predicts numerous new TFs active in neuronal and glial subtypes. Our data confirm the existence of several known cell types but also identify numerous new subtypes. Our analyses also reveal the downstream target genes and enriched TF binding motifs, complemented by ‘standard’ TF clustering.

### Neuronal birth order is correlated with metabolic state and spiking potential

We used our dataset to identify neuronal subtypes by their neurotransmitter type or neuroblast lineage. Both the analysis of diffusion components (Figure 2E) and SCENIC revealed a strong influence of the transcription factor *datilografo* (*dati*) (Figure 4A & Figure 5B/D). The enriched motif in this regulon is a peculiar AAAAAA motif that has been experimentally validated for Dati (FlyFactorSurvey database (Zhu et al., 2011)). The predicted target genes of Dati include *pros, chinmo, hb*, and *jim*. This finding prompted us to verify the expression of additional transcription factors that are related to neuronal birth order. These genes, mostly transcription factors and RNA binding proteins, are known to be specifically expressed during specific time windows of neuronal birth in the embryo and the larval brain (Maurange et al., 2008; Rossi et al., 2017). The activity of the Dati regulon, as well as the expression of *pros, mamo, br*, and *Imp*, represent a gradient orthogonally to the neurotransmitter cell types. Importantly, this gradient is not dependent on a specific genetic *Drosophila* strain as we identified it in both DGRP-551 and w^1118^ cells, and is not affected by ageing (Figure S7). We also checked additional birth-order factors like *chinmo* and *Syncrip* (Syp) (Liu et al., 2015; Zhu et al., 2006), but these are known to be regulated at the post-transcriptional level and do not show obvious changes at the mRNA level.

**Figure 5.**
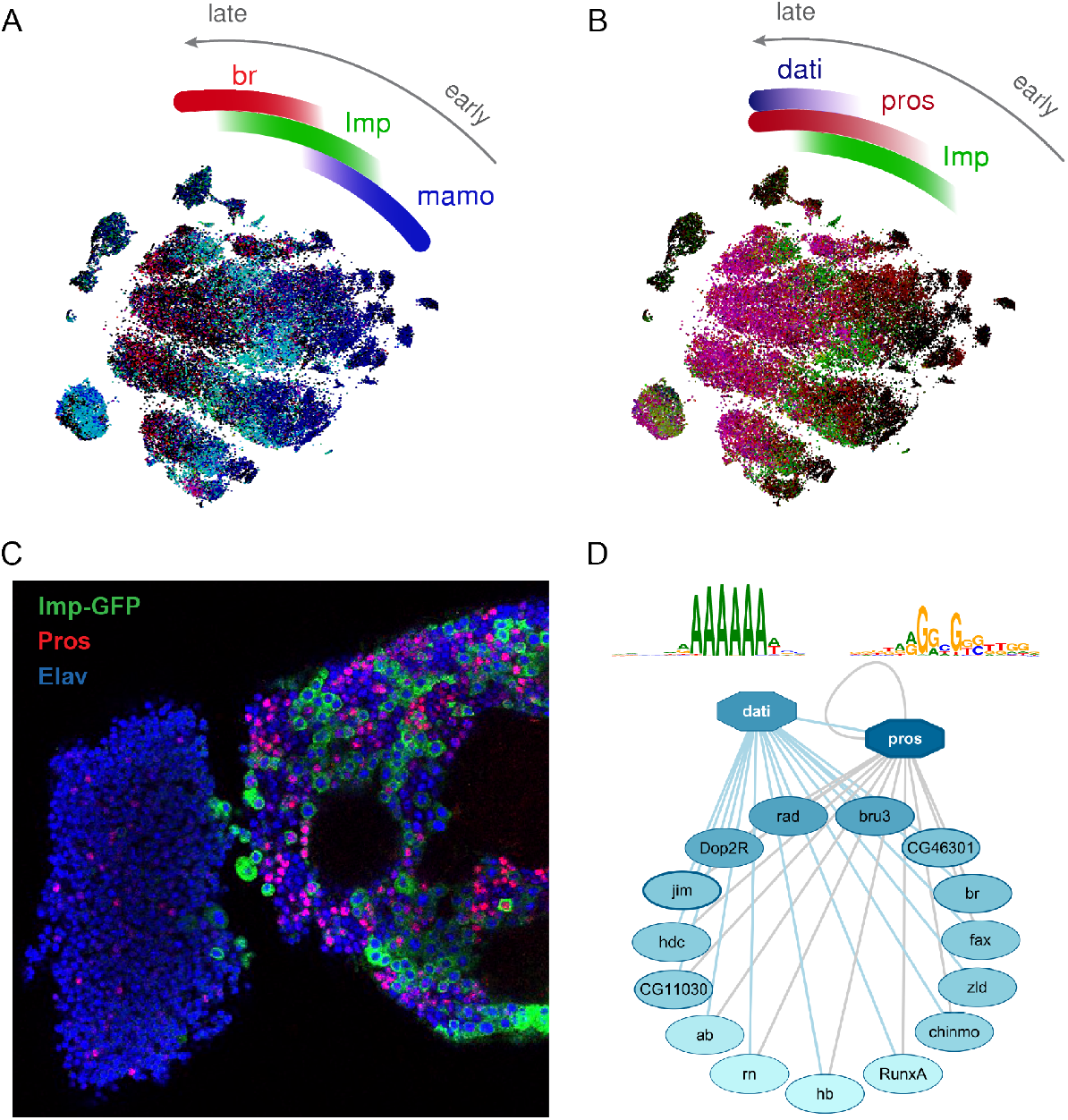
Regulatory state determined by birth-order factors. (A-B) Expression of early factors *Imp* and *mamo*, versus late factors like *broad, dati*, and *prospero*, represent a gradient in the SCENIC t-SNE from right (early) to left (late). (C) Immunostaining of Prospero and Imp-GFP showing a very strong disjoint expression of both factors, in agreement with the t-SNE. (D) Predicted Dati and Pros target genes in a co-regulated network, inferred by iRegulon through the significant enrichment of the Dati and Pros motifs (ranked 1^st^ and 2^nd^, respectively). The Dati motif is from FlyFactorSurvey and the Pros motif is the motif of the human homolog PROX1. See also Figure S7.

We validated expression of the Prospero protein using immunofluorescence. This confirmed that roughly half of the neurons in the central brain are Pros-positive and that the optic lobes are largely negative (Figure 5C). Similar to our scRNA-seq data, Pros expression appeared mutually exclusive with Imp. This immunofluorescence experiment also revealed no spatial bias in the central brain, with Prospero-positive cells being broadly distributed. These observations are in line with earlier observations of Dati immunofluorescence, which showed a similar “salt-and-pepper” labeling across the adult central brain (Schinaman et al., 2014). Interestingly, the ordering of neurons in the t-SNE plot correlated with their birth order, as illustrated by *Imp* and *mamo* being markers for early-born neurons, and *br, dati* and *pros* being markers for late-born neurons (Figure 5A/B). TF binding motif analysis using iRegulon not only confirmed the enrichment of the Dati motif, but also revealed significant enrichment (ranked 2^nd^) of the Prospero binding motif (Figure 5D), thus predicting a combinatorial gene regulatory network of Dati and Prospero to control this cell state.

Neurons positive and negative for *dati/pros* expression exist jointly in many of the neuronal subclusters (Figure 5B). To assess whether there are functional differences between the early-born *dati/pros*-negative neurons and the late-born *dati/pros*-positive neurons, we examined up-and down-regulated genes in early versus late-born neurons (early and late-born neurons were classified as having low and high expression of the Dati regulon, respectively). A rank-based Gene Ontology analysis using GOrilla (Eden et al., 2009) indicated that genes up-regulated in late-born neurons were most strongly enriched for “axon guidance” (GO:0007411; FDR 3.74x10^−15^), while genes up-regulated in early-born neurons were most strongly enriched for “electron transport chain” (GO:0022900; FDR 1.36x10^−56^). The latter prompted us to examine gene sets linked to a wider spectrum of metabolic processes (Figure 6A). Interestingly, this showed that early-born neurons have significantly higher expression of genes involved in oxidative phosphorylation and other mitochondria-related metabolic pathways (Tennessen et al., 2014). Following this, we compared the number of mitochondrial reads between early and late-born neurons. This confirmed a significantly higher number of such reads for early-born neurons, suggesting that these neurons may contain more active mitochondria (Kolmogorov-Smirnov p-value 2.22x10^−71^) (Figure 6B).

**Figure 6.**
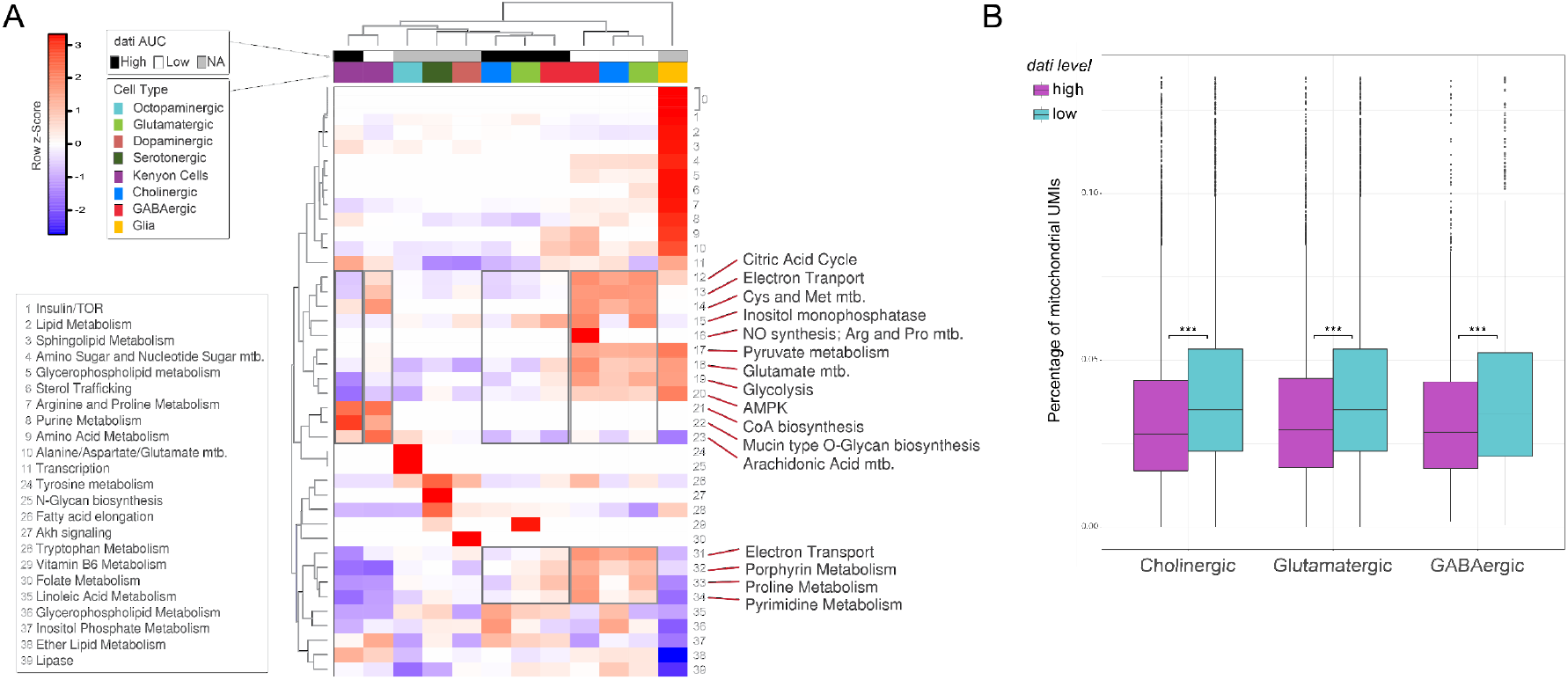
Dati expression correlates with metabolic state. (A) Aggregate expression of gene sets corresponding to metabolic pathways shows increased activity of oxidative phosphorylation in early-born (i.e. Dati-low) neurons. (B) Early-born (i.e. Dati-low) neurons have significantly more mitochondrial reads.

Based on this finding, we hypothesized that such metabolic activity could be related to neuronal spiking activities and/or intrinsic excitability, since initiation and propagating action potentials cause neurons to consume large amounts of energy (Attwell and Laughlin, 2001). In insect brains, some neurons have high action potential firing rates (high level of activity) (Flourakis et al., 2015), while other neurons are even non-spiking (Mu et al., 2012; Papadopoulou et al., 2011; Tootoonian et al., 2012). The activities and/or intrinsic excitability of different neurons could intuitively be in alignment with the *pros/dati* gradient. To test this hypothesis, we examined several classes of neurons and tested for correlations between *pros* expression and excitability or activity. Firstly, the Pdf-positive clock neurons are well-known as high-spiking neurons (Cao and Nitabach, 2008; Liu et al., 2014), and these are all *pros*-negative in our single-cell RNA-seq data (Figure 7A/B). We confirmed *in vivo* by anti-Pdf immunostaining that Pdf neurons show little Prospero immunoreactivity (Figure 7C). Secondly, a well-known class of non-spiking neurons are the antennal mechanosensory and motor center A1 neurons (AMMC-A1) (Tootoonian et al., 2012). Due to the lack of marker genes for this subtype, we could not localize these neurons in our dataset, but we could confirm *in vivo* that these neurons are Pros-positive by immunostaining (Figure 7D). Thirdly, we identified a small subset of 26 OPN neurons (out of the 543 OPN neurons; cfr. cluster 27 in Figure 3B) that are *pros*-negative. The lack of *pros* in these cells is likely not due to dropouts, because nearly all these (23 out of 26) cells are positive for *Imp*, which is anti-correlated with *pros* (Figure 7C). Fourthly, in the mushroom body we identify three subpopulations, which correspond to the alpha/beta, alpha’/beta’, and gamma lobes (Figure 7E). Of these, the alpha’/beta’ Kenyon cells, that are positive for DAT expression, are known to be most sensitive, and can be excited more easily than the other classes of Kenyon cells (Inada et al., 2017). In line with our expectation, this cluster had the lowest *pros* expression (Figure 7E). These examples indicate that *pros* expression levels negatively correlate with neuronal activity.

**Figure 7.**
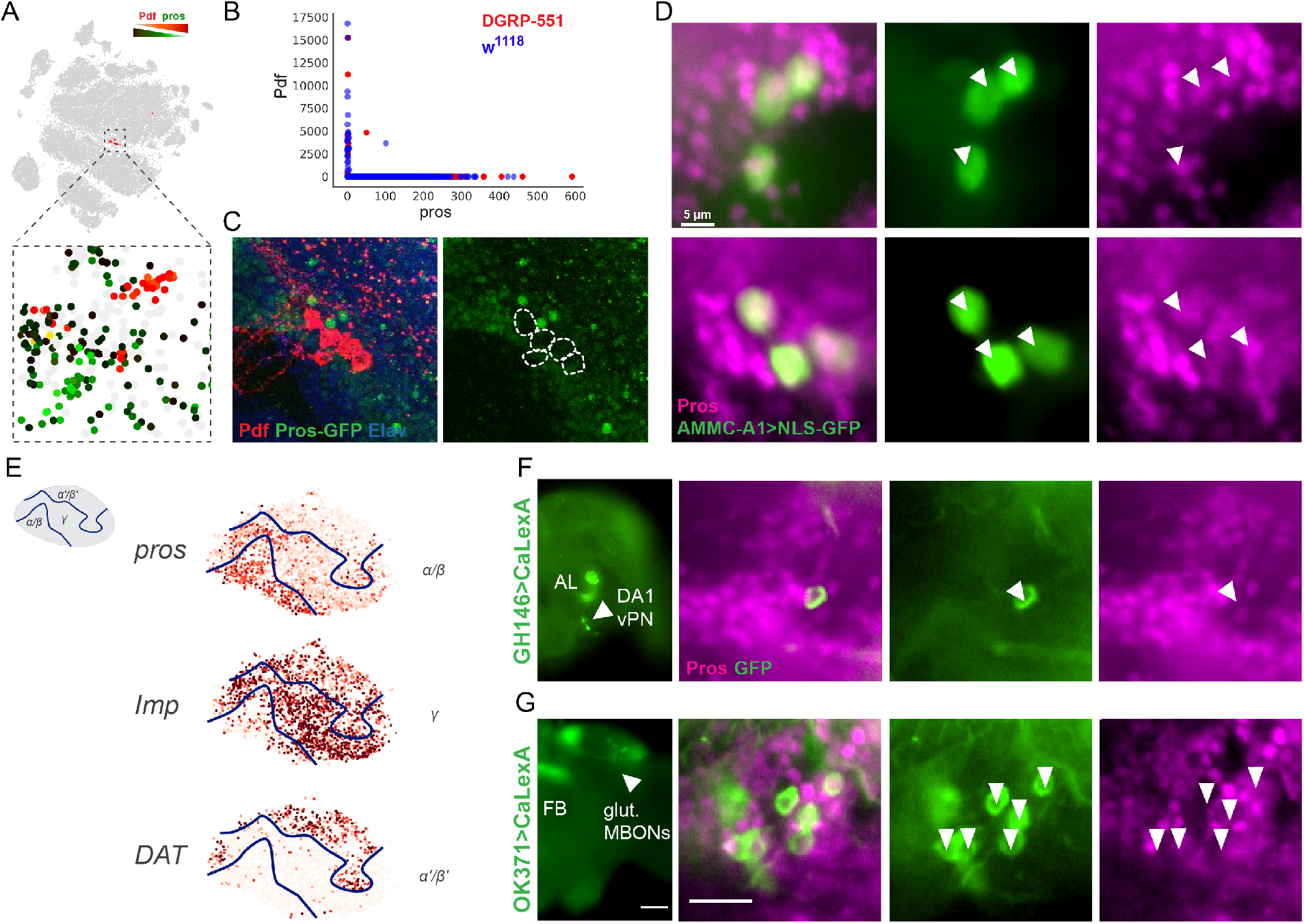
*pros* expression anticorrelates with neuronal excitability. (A-B) Spiking Pdf positive neurons are negative for *prospero*. (C) Confirmation by immunofluorescence that Pdf neurons are Pros-negative. (D) Nonspiking AMMC-A1 neurons are Pros-positive. (E) DAT positive mushroom body subcluster corresponding to alpha'/beta' Kenyon cells is most sensitive to excitation and *pros*-negative. (F) A small subset of OPNs is CaLexA-positive (GFP) and Pros-negative; these correspond to the DA1 vPN neurons involved in pheromone detection. (G) Glutamatergic MBONs are CaLexA-positive and Pros-negative.

We further tested the idea that Pros levels negatively correlate with neuronal activity and turned to an *in vivo* fluorescent activity reporter “UAS-CaLexA” (Masuyama et al., 2012). CaLexA labels active neurons using the calcium-dependent transcription factor NFAT to import LexA into the nucleus and activate a GFP reporter under the control of LexA binding sites. We expressed CaLexA in the OP-neurons, driven by the OPN-specific driver GH146-GAL4) and indeed identified a small set of OP-neurons that are GFP-positive but Pros-negative (Figure 7F). Based on their morphology and spatial localization, we were able to identify this subset of cells as the DA1 vPN neurons (Schlief and Wilson, 2007). These neurons convey signals for the male pheromone (cis-vaccenyl acetate or cVA). It is therefore plausible that they are more excitable than the rest of the OPN, which integrate multiple odors and need to keep noise levels low. Furthermore, we also performed a similar CaLexA experiment for the glutamatergic mushroom body output neurons (MBONs) (OK371-GAL4). Again, all GFP-positive neurons are Pros-negative (Figure 7G).

Together, these data are consistent with the idea that *pros/dati* negative neurons, with high metabolic activity, are more excitable neurons that are easier to generate spikes, whereas *pros/dati* positive neurons are less active/excitable neurons that may have a more integrative function, perhaps with more complex inputs, or with higher levels of plasticity. In conclusion, our work shows that all neurons in the fly brain are strongly influenced by a regulatory network of temporal transcription factors (e.g., *pros, dati, mamo, hb*) and RNA binding proteins (*Imp*), linking their birth order with their metabolic state and activity or excitability.

## Discussion

In this work, we report the first single-cell compendium of an entire intact complex brain. Single-cell sequencing technology is revolutionizing the identification of cell types, and as we show here, cellular states (ageing, regulatory networks, birth order and metabolic state). The brain in particular benefits enormously from the emergence of single-cell technologies, as only general cell types were known. We here identify numerous new subclusters and state changes and we define these either by transcription factor expression or by regulons. This work will greatly benefit efforts to build connectomes, since cellular identity is an important determinant of wiring programs.

Our work overcame a number of technical hurdles such as the small size of the fly neurons (1-5 micron cell diameter) and the lower RNA content in aged animals to ensure we sampled a sufficient amount of mRNA molecules per single brain cell. Previous single-cell datasets encompassed either the *Drosophila* embryo (Karaiskos et al., 2017) or smaller subsets of neurons (Zeisel et al., 2015; Lake et al., 2016; Tasic et al., 2016; Li et al., 2017). Our dataset now includes the single-cell transcriptomes for nearly all cells in the fly brain.

Our work is, to our knowledge, also the first to examine the ageing process at single-cell level in the brain. Ageing has long been associated with epigenetic changes (O’Sullivan and Karlseder, 2012), and an increased transcriptional heterogeneity (Martinez-Jimenez et al., 2017). However, it remains unclear whether there is a causal link between epigenetic and transcriptional erosion and if the observed increase in noise is due to changes in cellular composition of a tissue or whether there is rather an ageing mechanism common to all cell types.

In our data, we find a significant increase in transcriptional noise in older cells. Related work on single-cell ageing has been performed in human hematopoietic cells and pancreas, where increased transcriptional noise has similarly been described (Enge et al., 2017; Martinez-Jimenez et al., 2017). We furthermore link this increased transcriptional noise with an increased noise in nucleosome positioning. We speculate that this is due to lower amounts of histone proteins and transcription factors. Surprisingly, unsupervised clustering and trajectory analysis of neuronal transcriptomes do not yield strong age-dependent clustering. Instead, neurons of specific cell types cluster together, across all ages, indicating that cell fate is maintained in old cells, even though the total RNA content decreases and the noise increases. We furthermore conclude that ageing mechanisms are cell type-specific, since glial cells but not neurons show a clear ageing trajectory in their transcriptional profile.

Neuronal subtypes in *Drosophila* are strongly linked to three different axes. First, along their presynaptic specialization: cholinergic, GABAergic, glutamatergic, dopaminergic, peptidergic, octopaminergic and serotonergic clusters can be readily identified with different clustering approaches. Transcription factor combinations appear to underlie these clusters, in line with findings in *Caenorhabditis elegans* (Tursun et al., 2011). Second, a few dozen neuronal subtypes can be identified robustly across clustering approaches, mainly within cholinergic neurons in the optic lobe. Third, neurons in the central brain are differentiated by a combinatorial code of birth-order factor, such as *Imp, pros, dati, mamo*, and *br* (Maurange et al., 2008; Liu et al., 2015; Syed et al., 2017). Our results show that neuronal specialization occurring during development, through transcription factor combinations, is recapitulated by the expression patterns in the adult brain.

What emerged from the contrast between late-born *pros/dati*-positive and early-born *pros/dati*negative neurons are highly significant differences in metabolic pathways related to oxidative phosphorylation, suggesting more active mitochondria in *pros/dati*-negative neurons, throughout the entire central brain. We hypothesized that this could be related to neuronal activity (e.g., spiking versus non-spiking, increased electrical activity, higher calcium signaling, etc.), and confirmed that spiking neurons were generally *pros*-negative, while the non-spiking neurons we investigated were *pros*-positive. It is thus tempting to speculate that neurons are endowed with a particular metabolic potential when they are born, and that such a regulatory state may influence how neuronal networks are constructed during development, and how they function during life.

## Methods

### Fly husbandry and genotypes

Two genotypes of *Drosophila melanogaster* were used for the single-cell experiments in this study, namely DGRP-551 (Bloomington *Drosophila* Stock Center, BL#55026) from the *Drosophila* Genetics Reference Panel (Mackay et al., 2012) and w^1118^. For the ATAC-seq experiments, DGRP-820 (BL#25208), DGRP-907 (BL#28262), DGRP-913 (BL#28265) were used. The fly lines used in the immunofluorescence experiments are as follows: *y[1]w[*] Mi{PT-GFSTF.2}Imp[MI05901-GFSTF.2]* (Imp-GFP, BL#60237), *w[1118]; PBac{y[+mDint2]w[+mC]=pros-GFP.FPTB}VK00037* (pros-GFP, BL#66463), *w[*]; P{w[+mW.hs]=GawB}c767* (AMMC-A1-Gal4, BL#30848), *w[1118]; P{w[+mC]=UAS-GFP.nls}8* (UAS-NLS.GFP, BL#4776), *y[1] w[1118]; P{w[+mW.hs]=GawB}GH146* (GH146-Gal4, BL#30026). All flies were raised on a yeast based medium and kept at 25°C on a 12h/12h day/night light cycle. During ageing experiments, 20 females and 10 males were placed in each vial and flies were transferred to new vials every 4 days.

### Brain dissociation into single cells using collagenase I and dispase

Forty *D. melanogaster* adult brains (20 females and 20 males) were dissected and transferred to a tube containing 100 μL ice cold DPBS solution. After centrifugation at 800 rcf for 5 minutes, the supernatant was replaced by 50 μL of dispase (3 mg/mL, Sigma-Aldrich_D4818-2mg) and 75 μL collagenase I (100 mg/mL, Invitrogen_17100-017). Brains were dissociated at 25°C in a Thermoshaker (Grant Bio PCMT) for 2h at 25°C, 500 rpm. The enzymatic reaction was reinforced by pipette mixing every 15min. Cells were washed with 500 μL ice cold DPBS solution and resuspended in DPBS 0.01% BSA (for Drop-seq experiments) or DPBS 0.04% BSA (for 10x experiments). Cell suspensions were passed through a 10 μM pluriStrainer (ImTec Diagnostics_435001050). Cell viability and concentration were assessed by the LUNA-FL™ Dual Fluorescence Cell Counter.

### Drop-seq

We generated our own microfluidic devices starting from an acetate mask (Selba S.A.) through to the final PDMS devices as described previously (Macosko et al., 2015), utilizing a softlithobox installation (Blackhole Labs). Drop-seq was performed as previously described (Macosko et al., 2015), with the following modifications. During the second strand cDNA synthesis, beads were pooled in fractions of 4000 beads rather than 2000. After second strand synthesis, all fractions were pooled via column purification (Qiagen MinElute) and eluted in 12 μl, this pooled library was purified a second time using AMPure XP beads at 0.6x beads to sample and eluted in 12 μl. The purified library was quantified (Qubit) and 50 ng was used for library preparation using 1 μl of the Nextera Tn5 enzyme (Illumina) and incubating at 55’C for 5 minutes. Tagmented cDNA was purified using column purification (Qiagen MinElute), PCR was performed to add sequencing adapters and a final bead purification was performed (1.0x beads). The final purified library was quantified via qPCR and bioanalyzer.

### 10x Genomics

Single-cell libraries were generated using the GemCode Single-Cell Instrument and Single Cell 3’ Library & Gel Bead Kit v2 and Chip Kit (10x Genomics) according to the manufacturer’s protocol. Briefly, fly brain single cells were suspended in 0.4% BSA–PBS. About 8700 cells were added to each channel with a targeted cell recovery estimate of 5000 cells. After generation of nanoliter-scale Gel bead-in-EMulsions (GEMs), GEMs were reverse transcribed in a C1000 Touch Thermal Cycler (Bio Rad) programed at 53°C for 45 min, 85°C for 5 min, and hold at 4°C. After reverse transcription, single-cell droplets were broken and the single-strand cDNA was isolated and cleaned with Cleanup Mix containing DynaBeads (Thermo Fisher Scientific). cDNA was then amplified with a C1000 Touch Thermal Cycler programed at 98°C for 3 min, 12 cycles of (98°C for 15 s, 67°C for 20 s, 72°C for 1 min), 72°C for 1 min, and hold at 4°C. Subsequently, the amplified cDNA was fragmented, end-repaired, A-tailed and index adaptor ligated, with SPRIselect Reagent Kit (Beckman Coulter) with cleanup in between steps. Post-ligation product was amplified with a C1000 Touch Thermal Cycler programed at 98°C for 45 s, 14 cycles of (98°C for 20 s, 54°C for 30 s, 72°C for 20 s), 72°C for 1 min, and hold at 4°C. The sequencing-ready library was cleaned up with SPRIselect beads.

### Immunofluorescence

For immunofluorescence, young adult brains were dissected and processed as described in (Wang et al., 2002). The primary antibodies against mouse α-Prospero (1:20, #MR1A), rat α-Elav (1:100, #7E8A10) and, mouse α-Pdf (1:50, #PDF C7) were obtained from the Developmental Studies Hybridoma Bank. The antibody against rabbit GFP, (1:1000, #6455), the Alexa Fluor 647-488-555 secondary antibodies were obtained from Life Technologies.

### Whole tissue ATAC-seq

ATAC-seq was performed as previously described (Davie et al., 2015) with the only modification being the use of two adult brains (one male and one female) instead of 10 eye-antennal imaginal discs.

### ATAC-seq noise

ATAC-seq signal-to-noise ratios were calculated by dividing the aggregate peak height at the transcription start site of genes (TSS) by the aggregate peak height of the environment at 1000bp from the TSS.

### High-throughput sequencing

Before sequencing, the fragment size of every library was analyzed on a Bioanalyzer high sensitivity chip. The libraries were diluted to 2 nM and quantified by qPCR using primers against p5-p7 sequence. All 10x libraries were sequenced on a HiSeq4000 instrument (Illumina) with following sequencing parameters: 26 bp read 1 – 8 bp index 1 (i7) – 88 bp read 2. All Drop-seq libraries were sequenced on a NextSeq instrument (Illumina). Custom primers for index read 1 and read 2 were added to the Illumina sequencing primers. The following sequencing parameters were used: 51 bp read 1 – 6 bp index 1 (i7) – 8 bp index 2 (i5) – 26 bp read 2.

### Raw Datasets

The 10x fly brain samples were each processed (alignment, barcode assignment and UMI counting) with Cell Ranger (version 2.0.0) count pipeline. The Cell Ranger reference index was built upon the 3^rd^ 2017 FlyBase release (*D. melanogaster* r6.16) (Gramates et al., 2017).

### DGRP-551 & w^1118^ datasets

The “--recovered-cells” parameter was specified for each Cell Ranger run with the number of cells that we expect to recover for each individual single-library (Table S1, Table S2). Therefore, the number of cells in each sample was estimated by the intrinsic Cell Ranger 10x cell detection algorithm. The final raw DGRP-551 dataset was built by aggregating all the DGRP-551 samples using Cell Ranger aggregation pipeline without applying any Cell Ranger normalization method. The final raw w^1118^ dataset was built in the same way as the DGRP-551 dataset. Final dimensions of the raw DGRP-551 and w^1118^ datasets are 17,473 by 32,277, and 17,473 genes by 31,951 cells, respectively.

A custom method to select the cells, less stringent than the one from Cell Ranger, was used to build the large dataset of 129,236 cells (Figure S3C). A common technique to discriminate cell-associated barcodes and the barcodes associated with empty partitions is to plot the cumulative fraction of reads/UMIs for all barcodes and look at its “knee”. The single cells would be assigned to the barcodes below the cut-off defined by the location of the knee. A new method was used to detect this “knee”. This method is based on the 10x publicly available dataset 1:1 mixture of fresh frozen human (HEK293T) and mouse (NIH3T3) cells. The total UMIs associated to each raw barcode was computed per species and this was used to plot the cumulative fraction of UMIs plot for all barcodes. Each barcode is considered as a mixed cell if less than 95% of the UMIs are not associated to one of the 2 species. With this, the cumulative fraction of mixed cells was computed. This number is almost flat for first top barcodes but then starts to grow linearly. A linear model was fitted through the linear part of the cumulative fraction of mixed cells (25,000^th^ to 50,000^th^ barcode). The fitted model intersected the x-axis at ~12,419 which is very close to the expected number of cells (12k). A spline was fitted to the cumulative fraction of UMIs in order to find the tangent at this intercept. The angle of the tangent was then used as cut-off for each single-library dataset to discriminate the true cell barcodes from the empty partitions.

### Data Preprocessing

To clean the data, quality control and filtering based on Lun et al. (Lun et al., 2016) was applied on each dataset. based on 10x cell detection algorithm. Genes not expressed in any cells were removed, leaving 13,991 genes for the DGRP-551 dataset and 13,581 for the w^1118^ dataset. Then, a principal component analysis (PCA) based on quality metrics (calculated by the scater R-package (McCarthy et al., 2017b) was performed for each cell. If the cell’s metrics deviated more than 5 absolute deviations from the median, the cell was flagged as an outlier. The number of cells passing the quality controls are 29,137 and 27,765 respectively for DGRP-551 and w^1118^. Finally, low-abundance genes, defined as those with an average count below a filter threshold of 0.001 were removed. Dimensions of the final preprocessed DGRP-551 and w^1118^ datasets are 9,461 genes by 29,137 cells, and 9,349 genes by 27,765 cells, respectively.

### Seurat

The Seurat pipeline was executed on a combined dataset of DGRP-551 and ^w1118^ cells (Satija et al., 2015). First, cells were selected from the preprocessed files of each genotype and a matrix was constructed with all genes. Then the scater R-package was used to select genes as previously described. This matrix was then used as input. Next, the data were log-normalized with a scale factor of 10^4^. The latent variables, defined as the number of UMI and the percentage of mitochondrial reads per cell, were regressed out using a negative binomial model. The graph-based method from Seurat was used to cluster cells. The PCA was selected as dimensional reduction technique to use in construction of SNN graph. In order to select the number of PCs as input into the clustering algorithm, we performed a cross-validation step which lead us to select the first 82 PCs. We selected a resolution of 2.0, leading to 86 clusters. We identified markers specific to each cluster and calculated differential expression, using the default parameters and the “bimod” model.

### SCENIC

#### SCENIC fly databases

SCENIC was run as described in (Aibar et al., 2017). For this study, we created the motif databases that allow to use RcisTarget and SCENIC on *Drosophila* (dm6, gene names from FlyBase version FB2016_05). We scored the 20003 motifs (Position Weight Matrices) in v8 of the i-cisTarget (Imrichová et al., 2015) database on the regulatory regions up to 5KB upstream of each gene and within its introns. The gene-motif rankings were then built taking the best-scoring region for each motif and gene. These databases will be made available for download to use with RcisTarget.

#### DGRP-551 dataset

SCENIC was applied on the DGRP-551 dataset after removing genes that were not present in our dm6 motifs ranking database. The input UMI count matrix was normalized (CPM) and log-transformed (prior of 1). Targets genes that did not show a positive correlation based on the GENIE3 co-expression algorithm (> 0.03) in each TF-module were filtered out. After removing TF-modules having less than 20 genes, SCENIC found 3,064 TF-modules. A *cis*-regulatory motif analysis on each of the TF-modules with RcisTarget revealed 150 regulons. The top 1 percentile of the number of detected genes per cell was used to calculate the AUCell enrichment of each regulon in each cell. We performed a t-SNE clustering on the regulon AUC matrix to obtain a two-dimensional representation of the cell states.

#### w^1118^ dataset

The w^1118^ dataset was used to validate the regulons obtained from the DGRP-551 dataset. Precisely, all the filtered w^1118^ cells were scored through AUCell (Aibar et al., 2017) on the DGRP-551 regulons. A t-SNE clustering was then applied similarly as for DGRP-551 dataset.

#### DGRP-551 and w^1118^ combined and large datasets

All 56,902 cells originating from the two preprocessed dataset (DGRP-551 and w^1118^) were rescored on the regulons found in the DGRP-551 dataset using AUCell. The two-dimensional representation of the cell states for the combined dataset were computed using the t-SNE clustering algorithm on the regulon AUC matrix. This same method was applied for the large dataset (less stringent filtering) containing 129236 cells.

### Diffusion maps

Regulon AUC matrices were constructed for the main groups found in the SCENIC t-SNE, containing both DGRP-551 and w^1118^ cells. Next, diffusion components were calculated using the Destiny package (R) on these matrices (Angerer et al., 2016). The output was visualised using the seaborn package (Python). Regulons shown in the heatmaps were selected by taking the 9 regulons with the highest absolute correlation with the first diffusion component. Finally, the selected regulons were row normalized (row-based z-score).

### Metabolic pathways enrichment

Gene signatures of metabolic pathways were extracted from Tennessen et al. 2014 (Tennessen et al., 2014). We classified cells with a Dati regulon AUCell score higher than 0.3 as “Dati high” cells, and with a score less than 0.15 as “Dati low” cells. Each gene signature was then scored with AUCell. The resulting AUC matrix was row-normalized (row-based z-score).

### Pearson Distance

The Pearson distance was defined as 1-corr(x_ijk_,μ_ij_), with i being the replicate and j the cell type. This was calculated for all cells after which the cells were pooled into young (< 9 days) and old (> 9 days). Boxplots were generated using the seaborn package (Python), the t-test was performed using the scipy package (Python) and FDR (Benjamini & Hochberg) was calculated with the statsmodel package (Python).

### Gene Ontology enrichment analysis

To test for GO term enrichment, we used GOrilla (Eden et al., 2009).

## Acknowledgements

This work is funded by The Research Foundation - Flanders (FWO; grants G.0640.13 and G.0791.14 to S. Aerts), Special Research Fund (BOF) KU Leuven (grants PF/10/016 and OT/13/103 to S. Aerts), ERC Consolidator Grant (724226_cis-CONTROL to S. Aerts), and a “Opening the Future” grant to P.V. S. Aibar is supported by a PDM Postdoctoral Fellowship from the KU Leuven. K.D. is supported by a PhD fellowship from the agency for Innovation by Science and Technology (IWT). J.J. is supported by a PhD fellowship of The Research Foundation – Flanders (FWO). 10x Chromium was made available through VIB Tech Watch Funding. Computing was performed at the Vlaams Supercomputer Center (VSC). The funders had no role in study design, data collection and analysis, decision to publish or preparation of the manuscript. The authors thank Fernando Casares, Bassem Hassan, Dietmar Schmucker, Pierre Vanderhaeghen, Joris De Wit, and Sara De Sa Cesariny Calafate, for helpful discussions and for reviewing the manuscript.

## Author Contributions

Conceptualization, K.D., J.J. and S.Aerts; Software, K.D., S.Aibar, M.D.W. and G.H.; Validation, K.D., D.K., U.P., S.Aibar, S.M. and V.C.; Formal Analysis, K.D., J.J., D.K., U.P., S.Aibar, M.D.W., C.B.G.B., K.I.S., and S.Aerts; Investigation, K.D., J.J., D.K., U.P., S.M. and V.C.; Resources, B.V., J.L., S.L., and P.V.; Data Curation, K.D. and M.D.W.; Writing – Original Draft, K.D., J.J., D.K., U.P., S.Aibar, M.D.W., S.M., V.C., C.B.G.B., K.I.S., B.T., S.L., P.V. and S.Aerts; Visualization, K.D., J.J., D.K., U.P., S.Aibar, and M.D.W.; Supervision, K.D., and S.Aerts; Project Administration, K.D. and S.Aerts; Funding Acquisition, K.D., J.J. and S.Aerts

## References

Aibar, S., González-Blas, C.B., Moerman, T., Huynh-Thu, V.A., Imrichova, H., Hulselmans, G., Rambow, F., Marine, J.-C., Geurts, P., Aerts, J., et al. (2017). SCENIC: single-cell regulatory network inference and clustering. Nat. Methods 14, 1083–1086.

Angerer, P., Haghverdi, L., Büttner, M., Theis, F.J., Marr, C., and Buettner, F. (2016). destiny: diffusion maps for large-scale single-cell data in R. Bioinforma. Oxf. Engl. 32, 1241–1243.

Attwell, D., and Laughlin, S.B. (2001). An energy budget for signaling in the grey matter of the brain. J. Cereb. Blood Flow Metab. Off. J. Int. Soc. Cereb. Blood Flow Metab. 21, 1133–1145.

Buenrostro, J.D., Giresi, P.G., Zaba, L.C., Chang, H.Y., and Greenleaf, W.J. (2013). Transposition of native chromatin for fast and sensitive epigenomic profiling of open chromatin, DNA-binding proteins and nucleosome position. Nat. Methods 10, 1213–1218.

Cao, G., and Nitabach, M.N. (2008). Circadian control of membrane excitability in Drosophila melanogaster lateral ventral clock neurons. J. Neurosci. Off. J. Soc. Neurosci. 28, 6493–6501.

Chiang, A.-S., Lin, C.-Y., Chuang, C.-C., Chang, H.-M., Hsieh, C.-H., Yeh, C.-W., Shih, C.-T., Wu, J.-J., Wang, G.-T., Chen, Y.-C., et al. (2011). Three-Dimensional Reconstruction of Brain-wide Wiring Networks in Drosophila at Single-Cell Resolution. Curr. Biol. 21, 1–11.

Crocker, A., Guan, X.-J., Murphy, C.T., and Murthy, M. (2016). Cell-Type-Specific Transcriptome Analysis in the Drosophila Mushroom Body Reveals Memory-Related Changes in Gene Expression. Cell Rep. 15, 1580–1596.

Davie, K., Jacobs, J., Atkins, M., Potier, D., Christiaens, V., Halder, G., and Aerts, S. (2015). Discovery of transcription factors and regulatory regions driving in vivo tumor development by ATAC-seq and FAIRE-seq open chromatin profiling. PLoS Genet. 11, e1004994.

Eden, E., Navon, R., Steinfeld, I., Lipson, D., and Yakhini, Z. (2009). GOrilla: a tool for discovery and visualization of enriched GO terms in ranked gene lists. BMC Bioinformatics 10, 48.

Enge, M., Arda, H.E., Mignardi, M., Beausang, J., Bottino, R., Kim, S.K., and Quake, S.R. (2017). Single-Cell Analysis of Human Pancreas Reveals Transcriptional Signatures of Aging and Somatic Mutation Patterns. Cell 171, 321–330.e14.

Flourakis, M., Kula-Eversole, E., Hutchison, A.L., Han, T.H., Aranda, K., Moose, D.L., White, K.P., Dinner, A.R., Lear, B.C., Ren, D., et al. (2015). A Conserved Bicycle Model for Circadian Clock Control of Membrane Excitability. Cell 162, 836–848.

Gramates, L.S., Marygold, S.J., Santos, G.D., Urbano, J.-M., Antonazzo, G., Matthews, B.B., Rey, A.J., Tabone, C.J., Crosby, M.A., Emmert, D.B., et al. (2017). FlyBase at 25: looking to the future. Nucleic Acids Res. 45, D663–D671.

Hassan, B.A., Bermingham, N.A., He, Y., Sun, Y., Jan, Y.N., Zoghbi, H.Y., and Bellen, H.J. (2000). atonal regulates neurite arborization but does not act as a proneural gene in the Drosophila brain. Neuron 25, 549–561.

Imrichová, H., Hulselmans, G., Atak, Z.K., Potier, D., and Aerts, S. (2015). i-cisTarget 2015 update: generalized cis-regulatory enrichment analysis in human, mouse and fly. Nucleic Acids Res. 43, W57–64.

Inada, K., Tsuchimoto, Y., and Kazama, H. (2017). Origins of Cell-Type-Specific Olfactory Processing in the Drosophila Mushroom Body Circuit. Neuron 95, 357–367.e4.

Ito, M., Masuda, N., Shinomiya, K., Endo, K., and Ito, K. (2013). Systematic Analysis of Neural Projections Reveals Clonal Composition of the Drosophila Brain. Curr. Biol. 23, 644–655.

Janky, R., Verfaillie, A., Imrichová, H., Van de Sande, B., Standaert, L., Christiaens, V., Hulselmans, G., Herten, K., Naval Sanchez, M., Potier, D., et al. (2014). iRegulon: from a gene list to a gene regulatory network using large motif and track collections. PLoS Comput. Biol. 10, e1003731.

Karaiskos, N., Wahle, P., Alles, J., Boltengagen, A., Ayoub, S., Kipar, C., Kocks, C., Rajewsky, N., and Zinzen, R.P. (2017). The Drosophila embryo at single-cell transcriptome resolution. Science 358, 194–199.

Klaes, A., Menne, T., Stollewerk, A., Scholz, H., and Klämbt, C. (1994). The Ets transcription factors encoded by the Drosophila gene pointed direct glial cell differentiation in the embryonic CNS. Cell 78, 149–160.

Komiyama, T., Johnson, W.A., Luo, L., and Jefferis, G.S.X.E. (2003). From lineage to wiring specificity. POU domain transcription factors control precise connections of Drosophila olfactory projection neurons. Cell 112, 157–167.

Kremer, M.C., Jung, C., Batelli, S., Rubin, G.M., and Gaul, U. (2017). The glia of the adult Drosophila nervous system. Glia 65, 606–638.

Lake, B.B., Ai, R., Kaeser, G.E., Salathia, N.S., Yung, Y.C., Liu, R., Wildberg, A., Gao, D., Fung, H.-L., Chen, S., et al. (2016). Neuronal subtypes and diversity revealed by single-nucleus RNA sequencing of the human brain. Science 352, 1586–1590.

Li, H., Horns, F., Wu, B., Xie, Q., Li, J., Li, T., Luginbuhl, D.J., Quake, S.R., and Luo, L. (2017). Classifying Drosophila Olfactory Projection Neuron Subtypes by Single-Cell RNA Sequencing. Cell 171, 1206–1220.e22.

Li, X., Chen, Z., and Desplan, C. (2013a). Temporal Patterning of Neural Progenitors in Drosophila. Curr. Top. Dev. Biol. 105, 69–96.

Li, X., Erclik, T., Bertet, C., Chen, Z., Voutev, R., Venkatesh, S., Morante, J., Celik, A., and Desplan, C. (2013b). Temporal patterning of Drosophila medulla neuroblasts controls neural fates. Nature 498, 456–462.

Liu, S., Lamaze, A., Liu, Q., Tabuchi, M., Yang, Y., Fowler, M., Bharadwaj, R., Zhang, J., Bedont, J., Blackshaw, S., et al. (2014). WIDE AWAKE mediates the circadian timing of sleep onset. Neuron 82, 151–166.

Liu, Z., Yang, C.-P., Sugino, K., Fu, C.-C., Liu, L.-Y., Yao, X., Lee, L.P., and Lee, T. (2015). Opposing intrinsic temporal gradients guide neural stem cell production of varied neuronal fates. Science 350, 317–320.

Lun, A.T.L., McCarthy, D.J., and Marioni, J.C. (2016). A step-by-step workflow for low-level analysis of single-cell RNA-seq data with Bioconductor. F1000Research 5, 2122.

Mackay, T.F.C., Richards, S., Stone, E.A., Barbadilla, A., Ayroles, J.F., Zhu, D., Casillas, S., Han, Y., Magwire, M.M., Cridland, J.M., et al. (2012). The Drosophila melanogaster Genetic Reference Panel. Nature 482, 173–178.

Macosko, E.Z., Basu, A., Satija, R., Nemesh, J., Shekhar, K., Goldman, M., Tirosh, I., Bialas, A.R., Kamitaki, N., Martersteck, E.M., et al. (2015). Highly Parallel Genome-wide Expression Profiling of Individual Cells Using Nanoliter Droplets. Cell 161, 1202–1214.

Martinez-Jimenez, C.P., Eling, N., Chen, H.-C., Vallejos, C.A., Kolodziejczyk, A.A., Connor, F., Stojic, L., Rayner, T.F., Stubbington, M.J.T., Teichmann, S.A., et al. (2017). Aging increases cell-to-cell transcriptional variability upon immune stimulation. Science 355, 1433–1436.

Masuyama, K., Zhang, Y., Rao, Y., and Wang, J.W. (2012). Mapping neural circuits with activity-dependent nuclear import of a transcription factor. J. Neurogenet. 26, 89–102.

Maurange, C., Cheng, L., and Gould, A.P. (2008). Temporal transcription factors and their targets schedule the end of neural proliferation in Drosophila. Cell 133, 891–902.

McCarthy, D.J., Campbell, K.R., Lun, A.T.L., and Wills, Q.F. (2017a). Scater: preprocessing, quality control, normalization and visualization of single-cell RNA-seq data in R. Bioinforma. Oxf. Engl. 33, 1179–1186.

McCarthy, D.J., Campbell, K.R., Lun, A.T.L., and Wills, Q.F. (2017b). Scater: preprocessing, quality control, normalization and visualization of single-cell RNA-seq data in R. Bioinforma. Oxf. Engl. 33, 1179–1186.

Mu, L., Ito, K., Bacon, J.P., and Strausfeld, N.J. (2012). Optic glomeruli and their inputs in Drosophila share an organizational ground pattern with the antennal lobes. J. Neurosci. Off. J. Soc. Neurosci. 32, 6061–6071.

Omoto, J.J., Lovick, J.K., and Hartenstein, V. (2016). Origins of glial cell populations in the insect nervous system. Curr. Opin. Insect Sci. 18, 96–104.

O’Sullivan, R.J., and Karlseder, J. (2012). The great unravelling: chromatin as a modulator of the aging process. Trends Biochem. Sci. 37, 466–476.

Owald, D., Lin, S., and Waddell, S. (2015). Light, heat, action: neural control of fruit fly behaviour. Philos. Trans. R. Soc. Lond. B. Biol. Sci. 370, 20140211.

Papadopoulou, M., Cassenaer, S., Nowotny, T., and Laurent, G. (2011). Normalization for sparse encoding of odors by a wide-field interneuron. Science 332, 721–725.

Park, D., Veenstra, J.A., Park, J.H., and Taghert, P.H. (2008). Mapping peptidergic cells in Drosophila: where DIMM fits in. PloS One 3, e1896.

Robie, A.A., Hirokawa, J., Edwards, A.W., Umayam, L.A., Lee, A., Phillips, M.L., Card, G.M., Korff, W., Rubin, G.M., Simpson, J.H., et al. (2017). Mapping the Neural Substrates of Behavior. Cell 170, 393–406.e28.

Rossi, A.M., Fernandes, V.M., and Desplan, C. (2017). Timing temporal transitions during brain development. Curr. Opin. Neurobiol. 42, 84–92.

Satija, R., Farrell, J.A., Gennert, D., Schier, A.F., and Regev, A. (2015). Spatial reconstruction of single-cell gene expression data. Nat. Biotechnol. 33, 495–502.

Schinaman, J.M., Giesey, R.L., Mizutani, C.M., Lukacsovich, T., and Sousa-Neves, R. (2014). The KRÜPPEL-like transcription factor DATILÓGRAFO is required in specific cholinergic neurons for sexual receptivity in Drosophila females. PLoS Biol. 12, e1001964.

Schlief, M.L., and Wilson, R.I. (2007). Olfactory processing and behavior downstream from highly selective receptor neurons. Nat. Neurosci. 10, 623–630.

Syed, M.H., Mark, B., and Doe, C.Q. (2017). Steroid hormone induction of temporal gene expression in Drosophila brain neuroblasts generates neuronal and glial diversity. ELife 6.

Tahoe, N.M.A., Mokhtarzadeh, A., and Curtsinger, J.W. (2004). Age-related RNA decline in adult Drosophila melanogaster. J. Gerontol. A. Biol. Sci. Med. Sci. 59, B896–901.

Tasic, B., Menon, V., Nguyen, T.N., Kim, T.K., Jarsky, T., Yao, Z., Levi, B., Gray, L.T., Sorensen, S.A., Dolbeare, T., et al. (2016). Adult mouse cortical cell taxonomy revealed by single cell transcriptomics. Nat. Neurosci. 19, 335–346.

Tennessen, J.M., Bertagnolli, N.M., Evans, J., Sieber, M.H., Cox, J., and Thummel, C.S. (2014). Coordinated metabolic transitions during Drosophila embryogenesis and the onset of aerobic glycolysis. G3 Bethesda Md 4, 839–850.

Tootoonian, S., Coen, P., Kawai, R., and Murthy, M. (2012). Neural representations of courtship song in the Drosophila brain. J. Neurosci. Off. J. Soc. Neurosci. 32, 787–798.

Tulving, E., and Craik, F.I.M. (2005). The Oxford Handbook of Memory (Oxford, New York: Oxford University Press).

Tursun, B., Patel, T., Kratsios, P., and Hobert, O. (2011). Direct conversion of C. elegans germ cells into specific neuron types. Science 331, 304–308.

Wyss-Coray, T. (2016). Ageing, neurodegeneration and brain rejuvenation. Nature 539, 180–186.

Xiong, W.C., Okano, H., Patel, N.H., Blendy, J.A., and Montell, C. (1994). repo encodes a glial-specific homeo domain protein required in the Drosophila nervous system. Genes Dev. 8, 981–994.

Yardımcı, G.G., Frank, C.L., Crawford, G.E., and Ohler, U. (2014). Explicit DNase sequence bias modeling enables high-resolution transcription factor footprint detection. Nucleic Acids Res. 42, 11865–11878.

Zeisel, A., Muñoz-Manchado, A.B., Codeluppi, S., Lönnerberg, P., La Manno, G., Juréus, A., Marques, S., Munguba, H., He, L., Betsholtz, C., et al. (2015). Brain structure. Cell types in the mouse cortex and hippocampus revealed by single-cell RNA-seq. Science 347, 1138–1142.

Zhu, L.J., Christensen, R.G., Kazemian, M., Hull, C.J., Enuameh, M.S., Basciotta, M.D., Brasefield, J.A., Zhu, C., Asriyan, Y., Lapointe, D.S., et al. (2011). FlyFactorSurvey: a database of Drosophila transcription factor binding specificities determined using the bacterial one-hybrid system. Nucleic Acids Res. 39, D111–117.

Zhu, S., Lin, S., Kao, C.-F., Awasaki, T., Chiang, A.-S., and Lee, T. (2006). Gradients of the Drosophila Chinmo BTB-zinc finger protein govern neuronal temporal identity. Cell 127, 409–422.

